# Rescue of SARS-CoV-2 from a single bacterial artificial chromosome

**DOI:** 10.1101/2020.07.22.216358

**Authors:** Chengjin Ye, Kevin Chiem, Jun-Gyu Park, Fatai Oladunni, Roy Neal Platt, Tim Anderson, Fernando Almazan, Juan Carlos de la Torre, Luis Martinez-Sobrido

**Affiliations:** Texas Biomedical Research Institute, San Antonio, TX, USA; Department of Veterinary Microbiology, University of Ilorin, Nigeria; Department of Molecular and Cell Biology, Centro Nacional de Biotecnología (CNB-CSIC), Madrid, Spain; Department of Immunology and Microbiology, The Scripps Research Institute, La Jolla, California, USA

## Abstract

An infectious coronavirus disease 2019 (COVID-19) emerged in the city of Wuhan (China) in December 2019, causing a pandemic that has dramatically impacted public health and socioeconomic activities worldwide. A previously unknown coronavirus, Severe Acute Respiratory Syndrome CoV-2 (SARS-CoV-2), has been identified as the causative agent of COVID-19. To date, there are no United States (US) Food and Drug Administration (FDA)-approved vaccines or therapeutics available for the prevention or treatment of SARS-CoV-2 infection and/or associated COVID-19 disease, which has triggered a large influx of scientific efforts to develop countermeasures to control SARS-CoV-2 spread. To contribute to these efforts, we have developed an infectious cDNA clone of the SARS-CoV-2 USA-WA1/2020 strain based on the use of a bacterial artificial chromosome (BAC).

Recombinant (r)SARS-CoV-2 was readily rescued by transfection of the BAC into Vero E6 cells. Importantly, the BAC-derived rSARS-CoV-2 exhibited growth properties and plaque sizes in cultured cells comparable to those of the SARS-CoV-2 natural isolate. Likewise, rSARS-CoV-2 showed similar levels of replication to that of the natural isolate in nasal turbinates and lungs of infected golden Syrian hamsters. This is, to our knowledge, the first BAC based reverse genetics system for the generation of infectious rSARS-CoV-2 that displays similar features *in vivo* to that of a natural viral isolate. This SARS-CoV-2 BAC-based reverse genetics will facilitate studies addressing several important questions in the biology of SARS-CoV-2, as well as the identification of antivirals and development of vaccines for the treatment of SARS-CoV-2 infection and associated COVID-19 disease.

## INTRODUCTION

In December 2019, a previously unknown coronavirus (CoV) was isolated in Wuhan (China) from a patient with respiratory disease who had possible contact with wild animals (*1-3*). Since then, Severe Acute Respiratory Syndrome CoV-2 (SARS-CoV-2), the ethological agent responsible for coronavirus disease 2019 (COVID-19), has been detected in 216 countries, areas or territories, and it has been responsible for over 11,125,245 human cases and 528,204 deaths (https://www.who.int/emergencies/diseases/novel-coronavirus-2019). The unprecedented human health and socioeconomic impact of COVID-19 rivals only that of the “Spanish flu” pandemic, which occurred almost 100 years ago (*4-8*). To date, there are no United States (US) Food and Drug Administration (FDA)-approved prophylactics (vaccines) or specific therapeutics (antivirals) available for the prevention and treatment, respectively, of SARS-CoV-2-associated COVID-19 disease.

CoVs are enveloped, single-stranded, positive-sense RNA viruses belonging to the *Nidovirales* order, and responsible for causing seasonal mild-respiratory illness in humans (e.g. 229E, NL63, OC43, HKU1). However, two previous CoVs have been associated with severe illnesses and resulted in significant morbidity and mortality in humans. These include Severe Acute Respiratory Syndrome CoV (SARS-CoV) in 2002; and the Middle East Respiratory Syndrome CoV (MERS-CoV) in 2012 (*9*). Like the SARS-CoV, SARS-CoV-2 genome is approximately 30,000 bases in length. Nonetheless, a feature of SARS-CoV-2 that is unique among known betacoronaviruses, is the presence of a furin cleavage site in the viral spike (S) glycoprotein, a characteristic known to increase pathogenicity and transmissibility in other viruses (*10*).

The ability of generating recombinant viruses using reverse genetics approaches represents a powerful tool to answer important questions in the biology of viral infections. It will help us to understand the mechanisms of viral infection, transmission and pathogenesis, as well as to identify viral and host factors and interactions that control viral cell entry, replication, assembly and budding. In addition, reverse genetics facilitates the generation of recombinant viruses expressing reporter genes for their use in cell-based screening assays or *in vivo* models of infection for the rapid and easy identification of prophylactic and therapeutic approaches for the treatment of viral infections, as well as to generate attenuated forms of viruses for their implementation as safe, immunogenic, and protective live-attenuated vaccines (LAVs).

DNA plasmids that replicate in *E. coli* have been previously used for the cloning of many viral genomes and generation of reverse genetics systems. However, assembly of full-length cDNAs of viruses with a large vial genome in *E. coli* is very challenging technically due to toxicity or instability, or both, of sequences within the viral genome. Two recent papers have described the ability to assemble the full-length genome of SARS-CoV-2 by *in vitro* ligation (*11*) or homologous recombination in yeast (*12*) to overcome this problem. However, both approaches rely on production of the full-length viral genome RNA by *in vitro* transcription, a process that poses technical difficulties. Bacterial artificial chromosomes (BACs), which are maintained as a single copy in *E. coli*, have been previously described to establish reverse genetic system for large RNA viruses, including other CoVs (*13-15*). To date, the use of a single BAC for the rescue of recombinant (r)SARS-CoV-2 has not been yet described. In this article, we describe the development of a BAC-based reverse genetics system for the recovery of rSARS-CoV-2 from transfected Vero E6 cells. Importantly, our results show that rSARS-CoV-2 and the natural SARS-CoV-2 isolate have similar fitness in cultured cells and replicate to similar levels in a validated golden Syrian hamster model of SARS-CoV-2 infection and associated COVID-19 disease. This is the first description of a reverse genetics approach for the rescue of rSARS-CoV-2 based on the use of a single BAC. This approach will facilitate studies on the biology of SARS-CoV-2, as well as the identification and characterization of antivirals and the development of LAVs for the control of SARS-CoV-2 infection and associated COVID-19 disease.

## RESULTS

### Assembly of SARS-CoV-2 genome in the BAC

We used a BAC-based approach, similar to the one we previously described for Zika virus (ZIKV) (*16-21*) to assemble an infectious clone of SARS-CoV-2 based on the USA-WA1/2020 strain (**Fig. 1A**). We selected this SARS-CoV-2 strain because it was isolated from an oropharyngeal swab from a patient with respiratory illness in Washington, US. The viral sequence was deposited in PubMed, and the virus isolate was available from BEI Resources.

**Figure 1.**
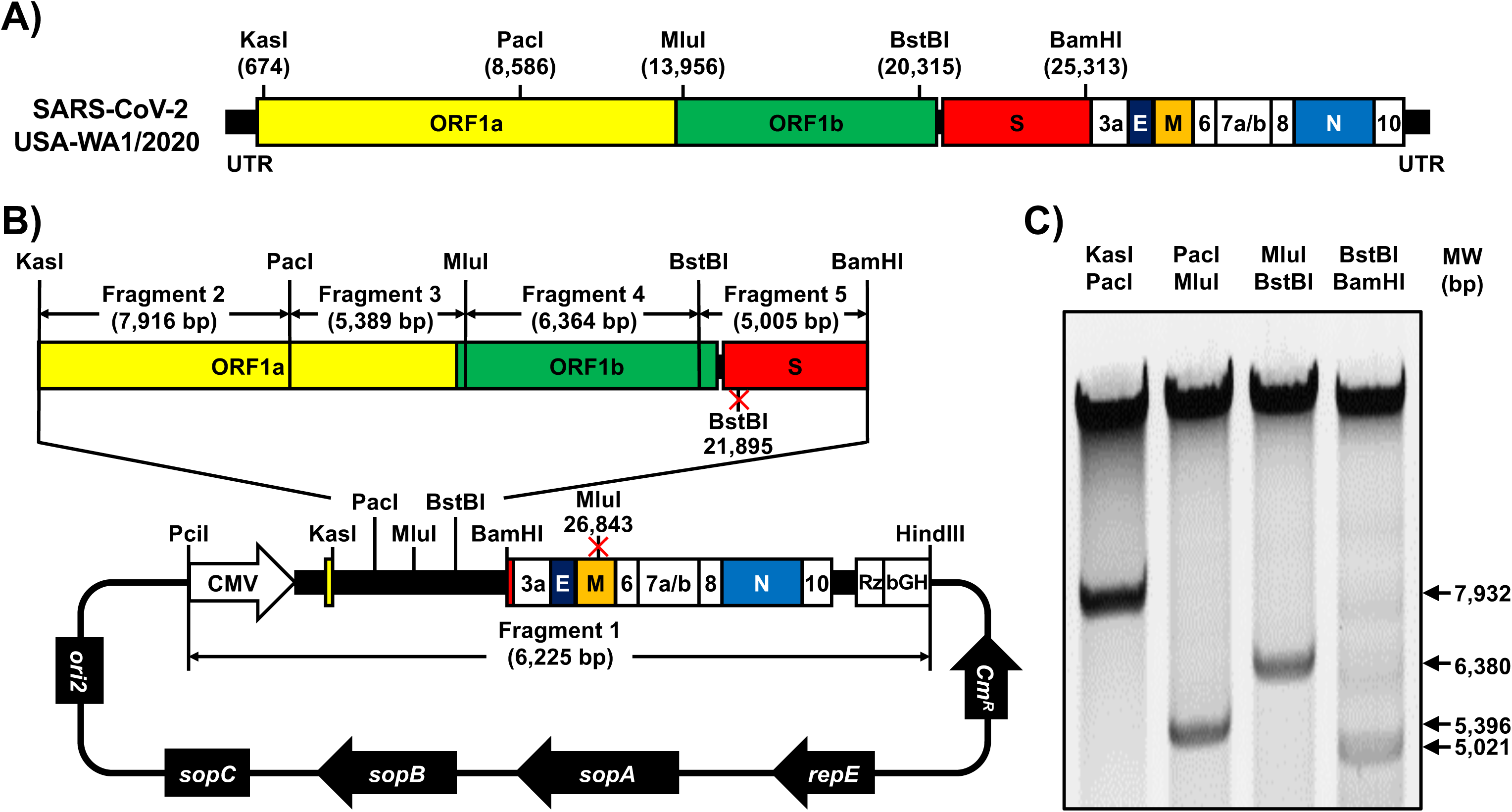
Assembly of SARS-CoV-2 genome into a BAC. **A**) **Schematic representation of SARS-CoV-2 genome:** The indicated restriction sites were used for cloning the entire viral genome (29,903 nucleotides) of SARS-CoV-2, USA-WA1/2020 strain, into the pBeloBAC11 plasmid. The open reading frames of the viral 1a, 1b, spike (S), envelop (E), matrix (M) and nucleocapsid (N) structural and the accessory (3a, 6, 7a, 7b, 8 and 10) proteins are also indicated. UTR, untranslated regions. Length is not in scale. **B-C**) **Assembly of the viral genome:** The full-length infectious cDNA clone was assembled by sequentially cloning the chemically synthesized fragments 1 to 5 covering the entire viral genome into the pBeloBAC11 plasmid, using the indicated restriction sites, under the control of the cytomegalovirus (CMV) promoter and flanked at the 3’ end by the Hepatitis Delta Virus (HDV) Ribozyme (Rz) and the bovine growth hormone (bGH) termination and polyadenylation sequences (**B**). The length of each of the chemically synthesized viral fragments is indicated. Ori2 indicates the origin of the replication of BAC. SopA, sopB and sopC are the elements to ensure each bacterial cell gets a copy of the BAC. Cm^R^ indicates chloramphenicol resistance. After assembly, the BAC clone harboring the entire viral genome was digested with the indicated restriction enzymes (top) and DNA products were analyzed in a 0.5% agarose gel (**C**).

We chemically synthesized the entire viral genome in 5 fragments that were assembled in the pBeloBAC plasmid using unique restriction enzymes and standard molecular biology approaches (**Fig. 1B**). After assembly of the 5 fragments, the BAC containing the entire viral genome was analyzed by restriction enzyme analysis (**Fig. 1C**). To facilitate the assembly of the viral genome and incorporate genetic tags to distinguish the rSARS-CoV-2 from the natural isolate, we introduced two silent mutations in the S (21,895 nt) and matrix, M (26,843 nt) viral genes that removed BstBI and MluI restriction sites, respectively (**Fig. 1B**).

### Rescue of rSARS-CoV-2

To recover rSARS-CoV-2, we used an experimental approach similar to that we previously described for mammarenaviruses (*22*) (**Fig. 2A**). Vero E6 cells were transfected with the SARS-CoV-2 BAC, or empty BAC as internal control, and were monitored for the presence of cytopathic effect (CPE), that were evident at 72 h post-transfection (**Fig. 2B**). Production of infectious virus (designated passage 0 [P0]) by transfected cells was 3.4×10^5^ PFU/ml (**Fig. 2C**). Recovery of rSARS-CoV-2 was confirmed by detection of viral antigen in fresh Vero E6 cells infected with tissue culture supernatants collected from SARS-CoV-2 BAC-transfected Vero E6 cells, but not from empty BAC-transfected Vero E6 cells, by immunofluorescence using a monoclonal antibody against the nucleocapsid (N) protein of SARS-CoV that cross-react with SARS-CoV-2 N (**Fig. 2D**).

**Figure 2.**
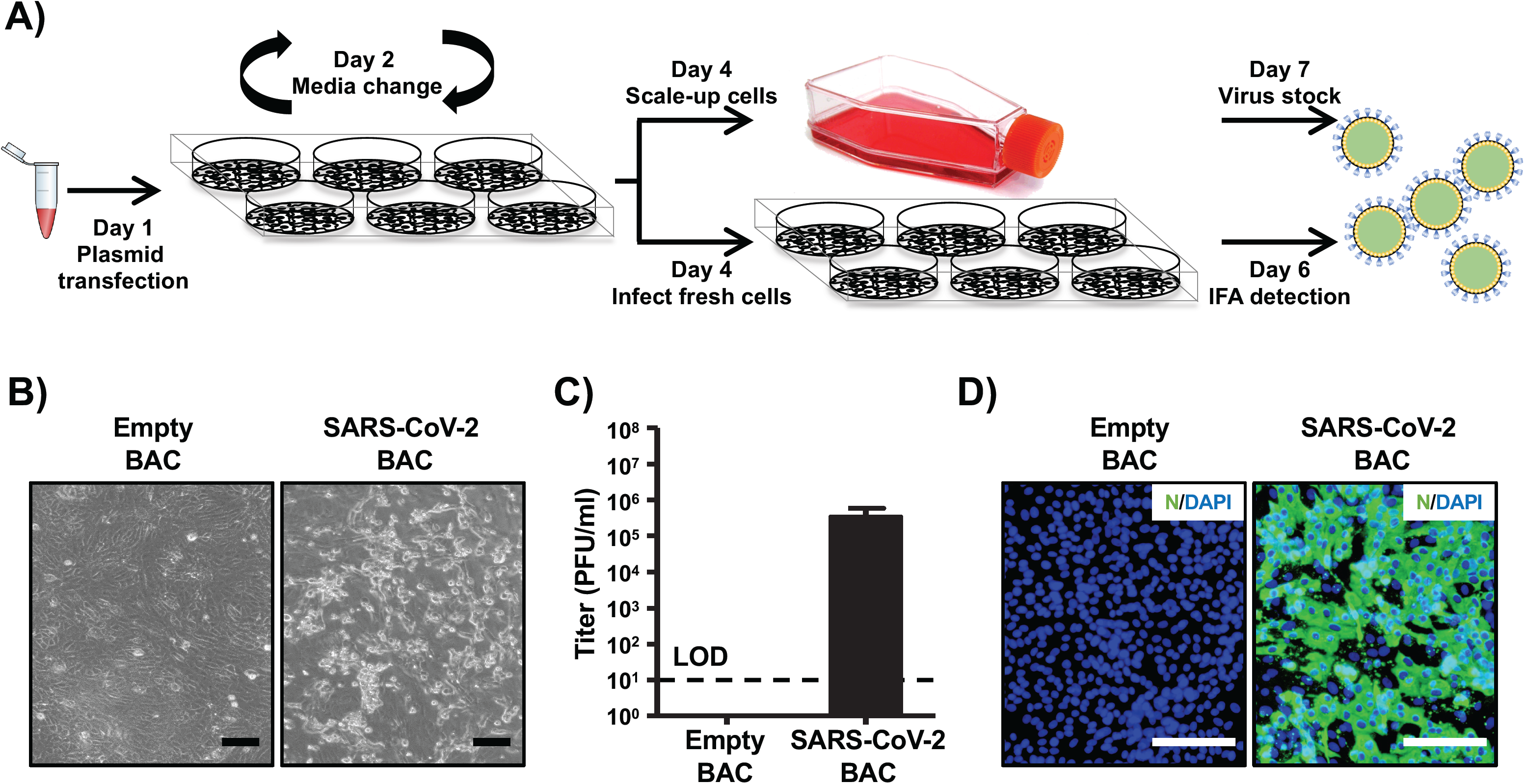
Rescue of rSARS-CoV-2. **A**) **Schematic representation of the approach followed to rescue rSARS-CoV-2:** Vero E6 cells (6-well plates, 10^6^ cells/well, triplicates) were transiently transfected with the SARS-CoV-2 BAC at day 1. After 24 h, transfection media was changed by post-infection media. At day 4, cells were split into T75 flask and the tissue culture supernatant was used to infect fresh Vero E6 cells (6-well plates, 10^6^ cells/well, triplicates). At 48 h post-infection, Vero E6 cells were fixed for detection of rSARS-CoV-2 by immunofluorescence and the tissue culture supernatant of the scaled-up Vero E6 cells was collected at 72 h. As internal control for this experiment, Vero E6 cells were transfected with the empty BAC. **B) CPE:** Images of empty or SARS-CoV-2 BACs transfected Vero E6 cells at 72 h post-transfection. Scale bars, 100 μm. **C) Viral titers:** Tissue culture supernatant from mock (empty BAC) or transfected Vero E6 cells in T75 flask was collected and titrated by immunofluorescence. Data were presented in mean ± SD. LOD: limit of detection. **D) IFA:** Vero E6 cells (6-well plate format, 10^6^ cells/well) infected with the tissue culture supernatants from transfected Vero E6 cells were fixed at 48 h post-infection and viral detection was carried out by using a SARS-CoV cross-reactive monoclonal antibody (1C7) against the N protein (green). Cellular nuclei were stained by DAPI (blue). Scale bars, 100 μm.

### Characterization of rSARS-CoV-2 *In vitro*

We first confirmed the genetic identity of the rescued rSARS-CoV-2. To that end, we used total RNA isolated from rSARS-CoV-2- and SARS-CoV-2-infected Vero E6 cells to amplified by RT-PCR a region in the M gene (nucleotides 26,488-27,784), in which a MluI restriction site was removed from the rSARS-CoV-2 cDNA via a silent mutation (**Fig. 1B**). As expected, the RT-PCR product from SARS-CoV-2-infected cells digested with MluI yielded two fragments with the size of 351 and 946 bp (**Fig. 3A**, bottom). In contrast, the RT-PCR product from rSARS-CoV-2-infected cells was not digested with MluI (**Fig. 3A**, bottom). We confirmed the mutation introduced in the MluI restriction site in the rSARS-CoV-2 by sanger sequencing (**Fig. 3B**).

**Figure 3.**
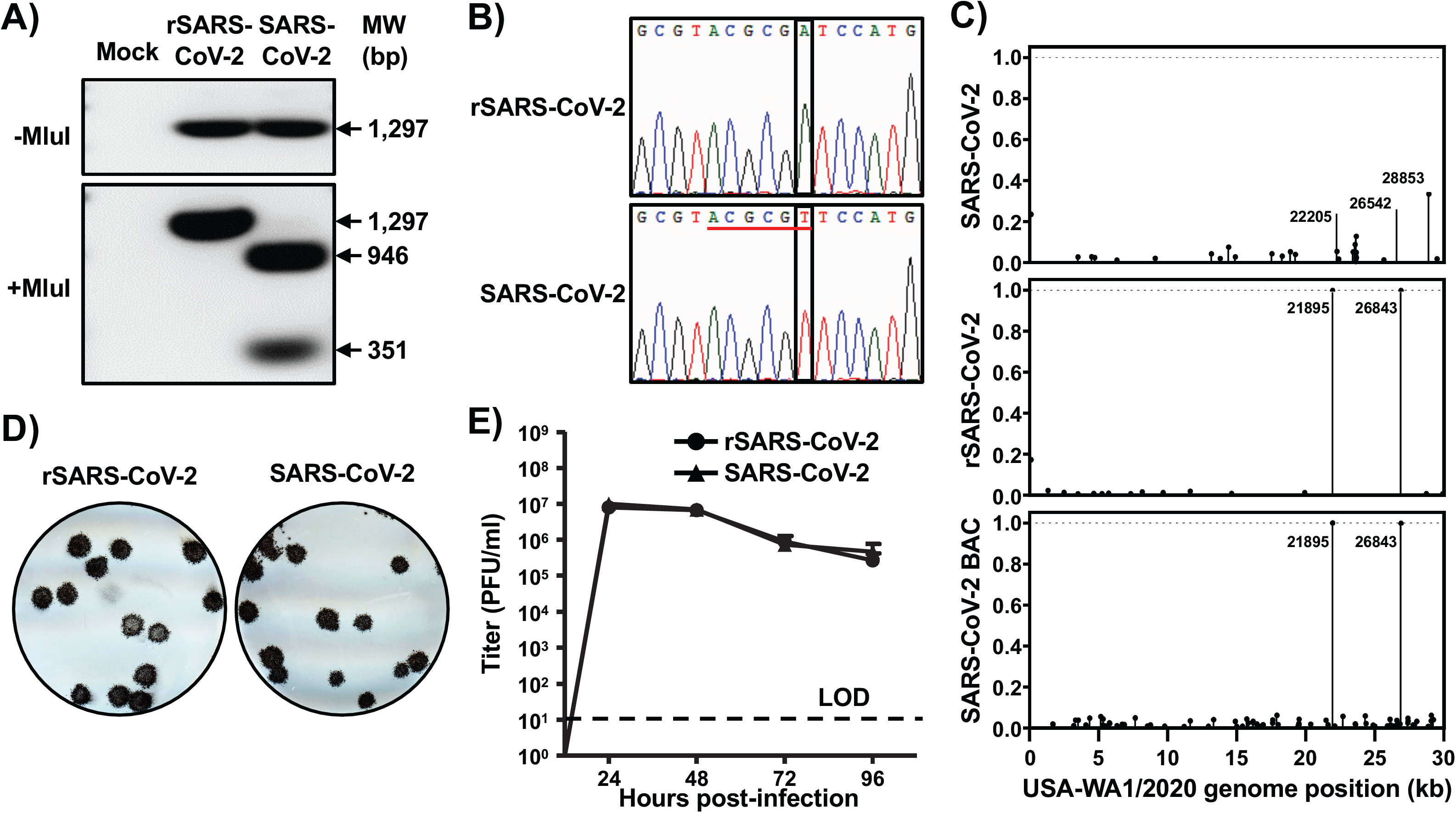
Characterization of rSARS2-CoV-2 *in vitro*. **A-B) Genotypic characterization:** Vero E6 cells (6-well plate, 10^6^ cells/well, triplicates) were mock-infected or infected (MOI 0.01) with rSARS-CoV-2 or the SARS-CoV-2 USA-WA1/2020 natural isolate. After 1 h viral adsorption at 37°C, Vero E6 cells were washed 3 times with PBS and incubated in post-infection media at 37°C. At 24 h post-infection, total RNA from Vero E6 cells was extracted and a 1,297 bp region of the M gene (nt 26,488 to 27,784) was amplified by RT-PCR. Amplified DNA was subjected to MluI digestion (**Figure 1**). Undigested (top) and digested (bottom) samples were separated in a 0.7% agarose gel (**A**). The RT-PCR amplified DNA product was also sequenced to verify the presence of the silent mutation in the MluI restriction site introduced in the viral genome of the rSARS-CoV-2 (**Figure 1**). The MluI restriction site is underlined in red and the silent mutation introduced to remove the MluI restriction site (T to A) is shown in the black box (**B**). **C**) **Verification of BAC and rSARS-CoV-2 sequences:** The SARS-CoV-2 non-reference allele frequency was calculated by comparing short reads to the reference genome of USA-WA1/2020 reference. All variants were at low frequency in P6 natural isolate (top), BAC (bottom), and rSARS-CoV-2 (middle) with the exception of introduced variants at positions 21,895 and 26,843, which were fixed in the BAC and rSARS-CoV-2 virus. Non-reference alleles present in less than 1% of reads are not shown. **D**) **Plaque phenotype:** Vero E6 cells (6-well plate, 10^6^ cells/well, triplicates) were infected with ∼20 PFU of rSARS-CoV-2 (left) or the natural SARS-CoV-2 isolate (right) for 1h at 37°C. After viral adsorption, cells were washed 3 times with PBS and overlaid with 2 ml of post-infection media containing 1% agar. After 72 h incubation at 37°C, cells were fixed and immunostained with the N protein 1C7 monoclonal antibody. **D**) **Growth kinetics:** Vero E6 cells (6-well plate, 10^6^ cells/well, triplicates) were infected (MOI 0.01) with rSARS-CoV-2 or the natural SARS-CoV-2 isolate for 1h at 37°C. After viral adsorption, cells were washed 3 times with PBS and incubated in 2 ml of post-infection media. At the indicated times post-infection, tissue culture supernatants were collected and viral titers were assessed by plaque assay (PFU/ml). Data were presented in mean ± SD. LOD: limit of detection.

To further characterize the genetic identity of rSARS-CoV-2, we used next generation sequencing (NGS) to determine the complete genome sequence of natural SARS-CoV-2 isolate from BEI Resources, and the rescue rSARS-CoV-2, as well as the BAC plasmid used to rescue rSARS-CoV-2. We examined 4.95M, 5.79M, and 5.44M reads for the natural virus isolated, BAC plasmid and rescued rSARS-CoV-2, resulting in coverages of 978x, 15,296x, and 1944x per sample, respectively. Introduced variants were not present in the SARS-CoV-2 but was effectively fixed in the BAC plasmid and rSARS-CoV-2. We confirmed the presence of the genetic markers at positions 21,895 (S) and 26,843 (M) in both, the BAC plasmid and rSARS-CoV-2 (allele frequencies > 99.9%). (**Fig. 3C**, middle and bottom).

Next, we compared rSARS-CoV-2 and the natural isolate of SARS-CoV-2 with respect their growth properties in Vero E6 cells. Both rSARS-CoV-2 and SARS-CoV-2 made uniform plaques with of similar size (**Fig. 3D**). Likewise, both rSARS-CoV-2 and SARS-CoV-2 exhibited similar growth kinetics and peak titers (**Fig. 3E**). These results confirmed the genetic identity of the rSARS-CoV-2 and its ability to replicate to the same extent than SARS-CoV-2 natural isolate in Vero E6 cells.

### Pathogenicity of rSARS-CoV-2 *in vivo*

Golden Syrian hamsters (Mesocricetus auratus) have been shown to be a good rodent animal model for investigating the replication, virulence and pathogenicity of both SARS-CoV (*23*) and SARS-CoV-2 (*24*) *in vivo*. To confirm that the rSARS-CoV-2 generated using the BAC-based reverse genetics exhibited the same replication capability, virulence and pathogenicity as the natural SARS-CoV-2 isolate *in vivo*, we infected golden Syrian hamsters intranasally with 2×10^4^ PFU of either rSARS-CoV-2 or the natural SARS-CoV-2 isolate. At days 2 and 4 post-infection, we collected the nasal turbinates (upper respiratory tract) and lungs (lower respiratory tract) from infected animals, as well as mock-infected controls, to assess the gross pathological changes (lungs) and the extent of viral replication (nasal turbinates and lungs). Mild multifocal congestion and consolidation were observed in 5-10% of the surface of lungs from rSARS-CoV-2 (**Fig. 4A, ii**) and SARS-CoV-2 (**Fig. 4A, iii**) infected animals at day 2 post-infection. As expected, the gross pathological lesions were more pronounced at day 4 post-infection, with severe multifocal to locally extensive congestion and consolidation (white arrows) in 40-50% of the surface of the lungs (**Fig. 4A, v** and vi). These lesions were widely distributed covering both the right (cranial, medial, and caudal lobes) and the left lobes of the lungs. Particularly, the presence of frothy exudate (black arrows) in the trachea of hamsters infected with either rSARS-CoV-2 or SARS-CoV-2 on day 4 post-infection indicates an ongoing bronchopneumonia. Of note, we did not observe significant differences in pathological lesion in the lungs at both days post-infection between animals infected with rSARS-CoV-2 or SARS-CoV-2 (**Fig. 4B**). Both rSARS-CoV-2 and SARS-CoV-2 replicated to similar levels in the lungs (**Fig. 4C**) and the nasal turbinates (**Fig. 4D**) of infected animals at days 2 and 4 post-infection, indicating that the genetically engineered rSARS-CoV-2 replicates to levels comparable to the natural isolate *in vivo*.

**Figure 4.**
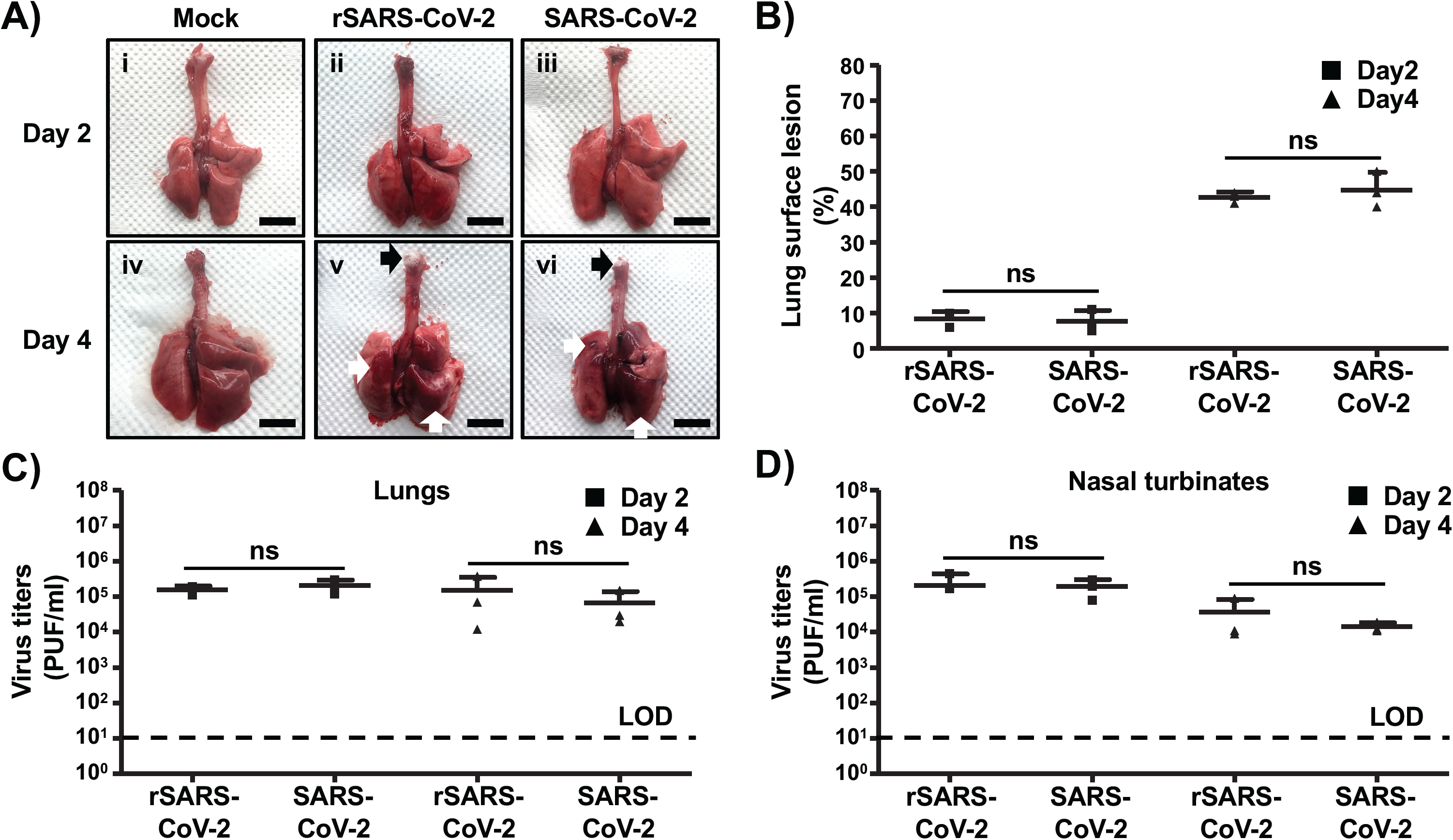
Pathogenicity of rescued rSARS-CoV-2 *in vivo*: Golden Syrian hamsters were mock-infected (n=2) or infected (n=6) with 2×10^4^ PFU of rSARS-CoV-2 or SARS-CoV-2. **A**) **Gross pathological lung lesions:** Animals were euthanized at 2 and 4 days post-infection, and lungs from mock-infected or infected (rSARS-CoV-2 and SARS-CoV-2) animals were observed for gross pathological changes, including congestion and atelectasis (white arrows) and frothy trachea exudates (black arrows). Scale bars, 1 cm. **B**) **Macroscopic pathology scoring analysis:** Distributions of pathological lesions, including consolidation, congestion, and atelectasis, were measured using ImageJ and represented as the percent of the total lung surface area (%). Ns, not significant. **C-D**) **Virus titers:** Viral titers in the lungs (C) and nasal turbinates (D) of rSARS-CoV-2 and SARS-CoV-2 infected golden Syrian hamsters were determined at days 2 and 4 post-infection (n=3/time point). Data represents the mean ± SD. Ns, not significant.

## DISCUSSION

The emergence of SARS-CoV-2, and associated COVID-19 disease, has created an unprecedented public health threat to humans (*25, 26*). To date, no FDA-approved prophylactics (vaccines) and/or therapeutics (antivirals) are available for the treatment of COVID-19, which has triggered a surge of scientific efforts to develop countermeasures to treat COVID-19 disease and control SARS-CoV-2 infection.

CoVs are single-stranded, positive-sense RNA viruses with one of the largest viral genomes (∼30 kilobases) among RNA viruses (*15*). Contrary to the situation with other single-stranded RNA viruses, the development of reverse genetics systems to recover recombinant CoV has been challenging due to toxicity or instability of viral cDNA in bacterial systems. To overcome this problem, several approaches have been used to successfully generate recombinant CoVs. This includes splitting of viral cDNA for propagation in different plasmids before assembly of a full-length genome by *in vitro* ligation (*27*), the use of homologous recombination in yeast (*28*), the insertion of linker sequences that disrupt the viral open reading frame and removed prior to transcription (*29*), the inactivation of cryptic promoters for bacterial RNA polymerase using silent mutagenesis (*30*), and the use of BACs (*13-15*).

In this study, we report the development of a full-length infectious clone of SARS-CoV-2 USA-WA1/2020 strain based on the use of a BAC. This is, to our knowledge, the first reverse genetics approach to generate rSARS-CoV-2 based on the transfection of a single BAC plasmid. The full-length cDNA copy of SARS-CoV-2 USA-WA1/2020 was sequentially assembled downstream of a cytomegalovirus (CMV) promoter into the pBeloBAC11 plasmid using synthetic fragments. After delivery of the BAC into host cells, the CMV promoter initiates the production of viral RNA from the nucleus of transfected cells by the cellular RNA polymerase II (Pol II). Although we described the generation of rSARS-CoV-2 from transfected Vero E6 cells, the use of the CMV promoter could also be applicable for generating rSARS-CoV-2 from other cell lines (*31*). Accordingly, we have been able to successfully rescue rSARS-CoV-2 using the BAC-based approach form human 293T and HeLa cells constitutively expressing the human angiotensin converting enzyme 2 (hACE2, data not shown).

The genetic identity of the rescued rSARS-CoV-2 was confirmed by sequencing. Notably, the rSARS-CoV-2 replicated in Vero E6 cells to levels comparable to the natural isolate as determined by growth kinetics and plaque assay. Importantly, using the golden Syrian hamster model of SARS-CoV-2 infection (*24*), we found that both rSARS-CoV-2 and the natural SARS-CoV-2 isolate have similar pathogenicity and growth capabilities in the upper and lower respiratory track of infected animals.

In summary, we have developed, for the first time, a powerful, reliable and convenient SARS-CoV-2 reverse genetics system based on the use of a BAC. The use of BAC-based reverse genetics for SARS-CoV-2 represents an excellent option to facilitate studies addressing a number of important concepts about the biology of SARS-CoV-2 infection. These include viral and host factors and interactions that control viral cell entry, replication, assembly and budding; the rescue of rSARS-CoV-2 with predetermined mutations in their genomes to examine their contribution to viral multiplication and pathogenesis; the develop of cell-based approaches to interrogate individual steps in the life cycle of SARS-CoV-2 to identify the mechanism of action of viral inhibitors; the generation of rSARS-CoV-2 expressing reporter genes for their use in cell-based screening assays or possibly *in vivo* models for the rapid and easy identification of viral inhibitors and/or neutralizing antibodies; and, the generation of rSARS-CoV-2 containing mutations in their viral genome that results in attenuation for their implementation as safe, immunogenic, stable and protective LAVs for the treatment of COVID-19 disease.

## MATERIAL AND METHODS

### Biosafety

All the *in vitro* and *in vivo* experiments with infectious SARS-CoV-2 were conducted under appropriated biosafety level (BSL) 3 and animal BSL3 (ABSL3) laboratories, respectively, at Texas Biomedical Research Institute (Texas Biomed). Experiments were approved by the Texas Biomed Institutional Biosafety (IBC) and Animal Care and Use (IACUC) committees.

### Cells and virus

African green monkey kidney epithelial cells (Vero E6, CRL-1586) were obtained from the American Type Culture Collection (ATCC, Bethesda, MD) and maintained in Dulbecco’s modified Eagle medium (DMEM) supplemented with 5% (v/v) fetal bovine serum, FBS (VWR) and 100 units/ml penicillin-streptomycin (Corning).

SARS-CoV-2 USA-WA1/2020 natural isolate was obtained from BEI Resources (NR-52281) and amplified on Vero E6 cells. This strain was selected because it was isolated from an oropharyngeal swab from a patient with respiratory illness in January 2020 in Washington, US. The SARS-CoV-2 USA-WA1/2020 sequence was available from Genbank (Accession No. MN985325).

### Sequencing

We generated short read sequencing libraries from the BAC and recovered SARS-CoV-2 viral RNA. For BAC sequencing, we followed the PCR-free KAPA HyperPlus Kit protocol, using 500 ng of input DNA. For SARS-CoV-2 viral RNA sequencing, we generated libraries using KAPA RNA HyperPrep Kit with a 45 min adapter ligation incubation including 6-cycle of PCR with 100 ng RNA and 7 mM adapter concentration. Samples were sequenced on an Illumina Hiseq X machine.

Raw reads were quality filtered using Trimmomatic v0.39 (*32*) and mapped to a SARS-CoV-2 reference genome (Genbank Accession No. MN985325) with Bowtie2 v2.4.1 (*33*). Genome coverage was quantified with MosDepth. V0.2.6 (*34*). We genotyped each sample for low frequency variants with LoFreq* v2.1.3.1 (*35*) and filtered sites with less than 100x read depth or minor allele frequencies less than 1%. Finally, we used SnpEff v4.3t (*36*) to identify the impact of potential variants on the protein coding regions in the SARS-CoV-2 reference genome.

### Assembly of the full-length of SARS-CoV-2 genome

Based on genomic information of SARS-CoV-2 USA-WA1/2020 isolate deposited in Genbank (Accession No. MN985325), the full-length genomic sequences were divided into 5 fragments (**Fig. 1**), synthesized *de novo* by Bio Basic (Ontario, Canada) and cloned into a high-copy pUC57 plasmid with designated restriction sites. A BstBI site in the S gene and a MluI site in the M gene were removed by silent mutation (**Fig. 1**). These mutations were introduced to ensure these restriction sites were unique and not present in the viral genome for the assembly of full-length SARS-CoV-2 genome and as molecular markers to distinguish the rescued rSARS-CoV-2 from the natural isolate (**Fig. 3**).

For the assembly of the entire viral genome in the BAC, fragment 1 was cloned into the pBeloBAC11 plasmid (NEB) linearized by PciI and HindIII digestion (**Fig. 1**). By using the preassigned restriction sites in fragment 1, the other 4 fragments were assembled sequentially by using standard molecular biology methods (**Fig. 1**). All the intermediate pBeloBAC11 plasmids were transformed into commercial DH10B electrocompetent *E. coli* cells (Thermo Fisher Scientific) using an electroporator (Bio-Rad) with the condition of 2.5kV, 600Ω and 10µF. The BAC containing the full-length SARS-CoV-2 genome, which had been analyzed by digestion using the restriction enzymes used to clone into the pBeloBAC 11 (**Fig. 1**), was also confirmed by deep sequencing.

### Rescue of rSARS-CoV-2

Virus rescue experiments were performed as previously described (*19*). Briefly, confluent monolayers of Vero E6 cells (10^6^ cells/well, 6-well plates, triplicates) were transfected, using LPF2000, with 4.0 μg/well of SARS-CoV-2 BAC, or empty BAC as internal control. After 24 h, transfection media was exchanged for post-infection media (DMEM supplemented with 2% (v/v) FBS), and cells were split and seeded into T75 flasks 48 h post-transfection. After incubation for another 72 h, tissue culture supernatants were collected, labeled as P0 and stored at -80°C. The P0 virus was used to infect fresh Vero E6 cells (10^6^ cells/well, 6-well plates, triplicates) (1 ml/well) for 48 h, and then cells were fixed and assessed for the presence of virus by immunofluorescence. After confirmation of the rescue, the P0 virus was subjected to 3 rounds of plaque purification a new virus stock (P3) was made and titrated for further *in vitro* and/or *in vivo* experiments.

### Immunofluorescence assay (IFA)

Vero E6 cells (10^6^ cells/well, 6-well plate format, triplicates) were mock-infected or infected (multiplicity of infection, MOI=0.01) with the natural USA-WA1/2020 isolate or rSARS-CoV-2. At 48 h post-infection, cells were fixed with 10% formaldehyde solution at 4°C overnight and permeabilized using 0.5% (v/v) Triton X-100 in PBS for 15 min at room temperature. Cells were incubated overnight with 1 μg/ml of a SARS-CoV cross-reactive N monoclonal antibody 1C7 at 4°C, washed with PBS, and stained with a FITC-labeled goat anti-mouse IgG (1:200). After washing with PBS, cells were visualized and imaged under a fluorescent microscope (Olympus).

### Plaque assay and immunostaining

Confluent monolayers of Vero E6 cells (10^6^ cells/well, 6-well plate format, triplicates) were infected with ∼20 PFU of SARS-CoV-2 USA-WA1/2020 or rSARS-CoV-2 for 1 h at 37°C. After viral adsorption, cells were overlaid with post-infection media containing 1% low melting agar and incubated at 37°C. At 72 h post-infection, cells were fixed overnight with 10% formaldehyde solution. For immunostaining, cells were permeabilized with 0.5% (v/v) Triton X-100 in PBS for 15 min at room temperature and immunostained using the N 1C7 monoclonal antibody (1 μg/ml) and the Vectastain ABC kit (Vector Laboratories), following the manufacturers’ instruction. After immunostaining, plates were scanned and photographed using a scanner (EPSON).

### Virus growth kinetics

Confluent monolayers of Vero E6 cells (10^6^ cells/well, 6-well plate format, triplicates) were infected (MOI=0.01) with SARS-CoV-2 USA-WA1/2020 or rSARS-CoV-2. After 1 h virus adsorption at 37°C, cells were washed with PBS and incubated in post-infection media at 37°C. At the indicated times after infection, viral titers in tissue culture supernatants were determined by plaque assay and immunostaining using the N monoclonal antibody 1C7, as previously described (*37*).

### RNA extraction and RT-PCR

Total RNA from SARS-CoV-2 USA-WA1/2020 or rSARS-CoV-2 infected (MOI=0.01) Vero E6 cells (10^6^ cells/well, 6-well plate format) was extracted with TRIzol Reagent (Thermo Fisher Scientific) according to the manufacturer’s instructions. RT-PCR amplification of the viral genome spanning nucleotides 26,488-27,784, was performed using Super Script II Reverse transcriptase (Thermo Fisher Scientific) and Expanded High Fidelity PCR System (Sigma Aldrich). The amplified 1,297 RT-PCR products were digested with MluI (NEB). Amplified DNA products, undigested or digested with MluI were subjected to 0.7% agarose gel analysis. Gel-purified PCR fragments were subjected to sanger sequencing (ACGT). All primer sequences used for RT-PCR are available on request.

### Pathogenicity studies in golden Syrian hamsters

Twelve-week-old female golden Syrian hamsters were purchased from Charles River and maintained in the animal facility at Texas Biomed under specific pathogen-free conditions. Golden Syrian hamsters were infected (1.0 ×10^4^ PFU) intranasally with either the rSARS-CoV-2 or the natural USA-WA1/2020 isolate in a final volume of 100 µl following gaseous sedation in an isoflurane chamber. After viral infection, hamsters were humanely euthanized on days 2 and 4 post-infection to collect nasal turbinates and lungs.

### Measurement of viral loads in nasal turbinates and lungs

Nasal turbinates and lungs from mock-, SARS-CoV-2- and SARS-CoV-2-infected golden Syrian hamsters were homogenized in 2 ml of PBS for 20 s at 7,000 rpm using a Precellys tissue homogenizer (Bertin Instruments). Tissue homogenates were centrifuged at 12,000 g (4 °C) for 5 min, and supernatants were collected for the measurement of viral loads. Confluent monolayers of Vero E6 cells (96-plate format, 4 × 10^4^ cells/well, duplicate) were infected with 10-fold serial dilutions of the supernatants from the tissue homogenates. After viral adsorption for 1 h at 37 °C, cells were washed 3 times with PBS before adding fresh post-infection media containing 1% microcrystalline cellulose (Avicel, Sigma Aldrich). Cells were further incubated at 37 °C for 24 h. Plates were then inactivated in 10% neutral buffered formalin (Thermo Fisher Scientific) for 24 h. For immunostaining, cells were washed three times with PBS and permeabilized with 0.5% Triton X-100 for 10 min at room temperature. Then, cells were blocked with 2.5% bovine serum albumin (BSA) in PBS for 1 h at 37 °C, followed by incubation with 1 µg/ml of the anti-N SARS-CoV monoclonal antibody 1C7 diluted in 1% BSA for 1 h at 37 °C. After incubation with the primary antibody, cells were washed three times with PBS, counterstained with the Vectastain ABC kit, and developed using the DAB Peroxidase Substrate kit (Vector Laboratory, Inc, CA, USA) according to the manufacturers’ instructions. Virus titers are indicated as PFU/ml.

### Evaluation of lung pathological lesions

Macroscopic pathology scoring was evaluated using ImageJ software to determine the percent of the total surface area of the lung (dorsal and ventral view) affected by consolidation, congestion, and atelectasis, as previously described (*38*).

### Statistical analysis

Data representative of three independent experiments in triplicates have been used. All data represent the mean ± standard deviation (SD) for each group and analyzed by SPSS13.0 (IBM). A two-tailed Student’s t-test was used to compare the mean between two groups. *P* values less than 0.05 (*P*<0.05) were considered statistically significant.

## ACKNOWLEDGEMENTS

We want to thank Dr. Thomas Moran at the Icahn School of Medicine at Mount Sinai for providing us with the SARS-CoV cross-reactive N monoclonal antibody 1C7. We also want to thank BEI Resources for providing the SARS-CoV-2 USA-WA1/2020 isolate (NR-52281) and Marina McDew-White and Robbie Diaz for constructing the NGS libraries. Finally, we would also like to thank members at our institutes for their efforts in keeping them fully operational during the COVID-19 pandemic and the BSC and IACUC committees for reviewing our protocols in a time efficient manner. We would like to dedicate this manuscript to all COVID-19 victims and to all heroes battling this disease.

